# Perceptual choice and motor signals in mouse somatosensory cortex

**DOI:** 10.1101/2024.12.06.627272

**Authors:** Genki Minamisawa, Mingyuan Dong, Daniel H. O’Connor

## Abstract

Somatosensory cortex activity relates both to sensation and movement, reflecting their intimate relationship, but the extent and nature of sensory-motor interactions in the somatosensory cortex remain unclear. Here, we investigated perception-related sensory and motor signals in the whisker areas of mouse primary (wS1) and secondary (wS2) somatosensory cortices. We recorded neuronal activity while mice performed a whisker detection task using two alternative lickports, one each to indicate the presence or absence of a whisker deflection on a given trial. One group of mice reported the presence of the whisker stimulus by licking at the port on the same (“congruent”) side of the animal as the stimulated whisker, whereas a second group of mice did so by licking at the opposite (“incongruent”) side. Activity of single neurons in wS1 and wS2 correlated with perceptual choice. This choice-related activity was enhanced when responding to the congruent side. wS2 neurons projecting along two output pathways—to wS1 or to whisker secondary motor cortex, wM2—also showed choice-related activity, but differed in their dependence on congruence and in the effects of optogenetic manipulation. Thus, somatosensory cortex contains pathway- and action-specific choice-related activity.

## INTRODUCTION

The sensory cortex activity that underlies tactile detection has been the focus of numerous studies in rodents using Go/NoGo task designs in which animals are trained to lick in response to a whisker deflection stimulus^1–4^. In these tasks, animals are typically trained to lick at a water spout to indicate the presence of a whisker stimulus on a given trial (a Go response), or to withhold licking in the absence of the stimulus (a NoGo response). A consistent finding is that, after brief deflections of a single whisker, neurons in the whisker regions of primary (wS1) and secondary (wS2) somatosensory cortices show higher levels of activity on trials when animals correctly lick to report the presence of the whisker deflection compared with when they failed to report the stimulus by not licking ^1–4^. Activity in wS1 and wS2 can predict the licking associated with tactile detection on a trial-by-trial basis^1,3^. This detection-related activity is transmitted bidirectionally along pathways linking wS1 and wS2^1^.

Optogenetically silencing wS1 or wS2, in order to inhibit the cortical response to the whisker stimulus, causes reductions in the detection performance of mice, indicating a role for wS1/wS2 in tactile detection tasks. Moreover, detection-related activity occurring even in time windows “late” (>100 ms) following the stimulus onset can impact task performance, as performance is impaired by wS1 silencing specifically during this period^2^. Together, this prior work has established that post-stimulus activity in wS1 and wS2 is not only evoked by whisker deflection, but is also associated with the Go/NoGo licking response.

In Go/NoGo tasks, however, it is not clear to what extent late activity in wS1 and wS2 reflects the perceptual event of stimulus detection, motor-related activity associated with a lick response, or some combination of sensory and motor signals. Defining the contributions of these different signals to activity in the whisker sensorimotor processing stream is an important area of current research^5^.

Here, we made single-unit recordings in wS1 and wS2, together with optogenetic perturbations, while mice performed a tactile detection task using directional licking at one of two ports to report either the presence or the absence of a brief whisker deflection on each trial. Mice made a lick response whether they detected the stimulus or not, allowing us to identify cortical activity associated with perceptual choice rather than licking per se. This choice-related activity was prominent among wS2→wS1 neurons and was enhanced when the direction of licking and the side of the stimulus were congruent. Optogenetic excitation of wS2→wS1 neurons promoted detection more strongly for congruent responses.

## RESULTS

### Distinct choice- and licking-related activity in wS1 and wS2

To separate perceptual choice-related activity from activity related to licking per se, we trained head-fixed mice to use licking to report on each trial whether they detected or did not detect a tactile stimulus (Figure 1A). On a random half of the trials, 0.5 s after an auditory cue a brief (25 ms) deflection was applied to a single whisker on the right side of the face (“Stim” trials; Figure 1B). On the other half of the trials, the stimulus was omitted (“NoStim” trials). Mice were trained to report the presence or absence of the stimulus on each trial by licking at one of two possible lickports, positioned symmetrically on either side of the mouse’s midline. An initial cohort of mice (n = 14) was trained to indicate the presence of the stimulus by licking at the lickport positioned to the right side of the midline, and the absence of the stimulus by licking to the lickport positioned on the left side (Right-stim→Right-lick contingency; Figure 1A). We refer to the ports used to indicate stimulus presence and absence as the “Stim port” and the “NoStim port”, respectively.

**Figure 1.**
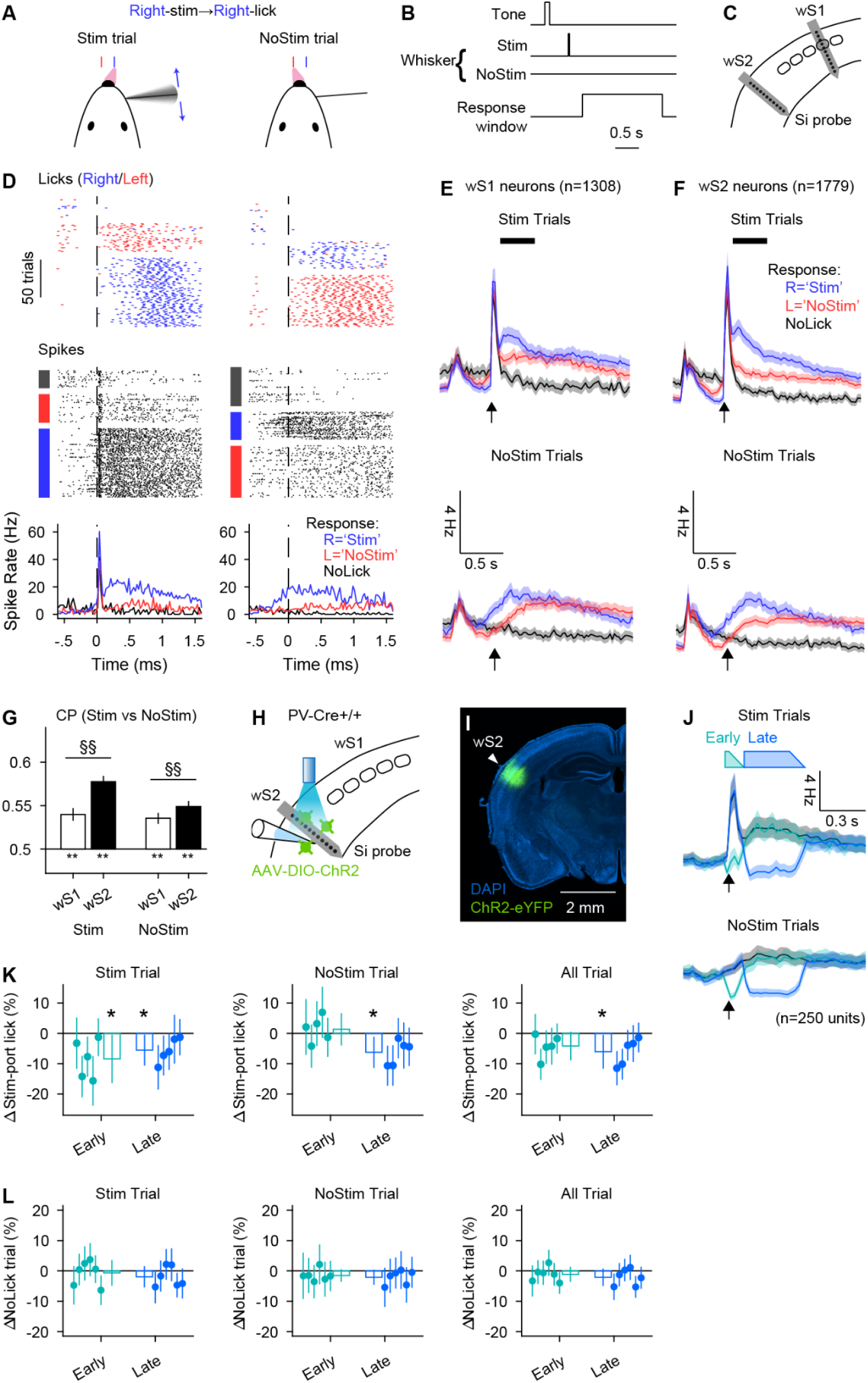
Choice- and licking-related activity in wS1 and wS2. (**A**) Whisker detection task with Right-stim→Right-lick (congruent) contingency. (**B**)Time course of the task. (**C**) Silicon probes were acutely implanted to wS1 and wS2 during the task. (**D**) Activity of an example neuron in a session. From top to bottom, lick and spike raster plots sorted by trial outcomes, and spike rate histogram triggered by the expected stimulus onset. Left and right columns correspond to data from Stim and NoStim trials, respectively. Lick directions were indicated by colors of the ticks or boxes on the left (Left: red, Right: blue). (**E**) Spike rate of all silicon probe-recorded wS1 neurons during Stim and NoStim trials. Expected stimulus onsets are indicated by black arrows. Shadings are 95% confidence intervals. Black box above the spike rate plot indicates the analysis window ([0, 500 ms] from the expected stimulus time) for choice probability in panel G. 31 recordings from 13 animals. (**F**) Same as E for wS2 neurons. 38 recordings from 14 animals. (**G**) Choice probabilities of Stim- over NoStim-port licking. *, **: p<.05 and <.01, Student’s t-test (null hypothesis: average CP=0); §, §§: p<.05 and 0.01, Two-sample t-test (null hypothesis: CP_S1_ = CP_S2_). (**H**-**I**) wS2 was inhibited by photo-excitation of PV-positive inhibitory interneurons. (**J**) Spike rates of wS2 neurons that were not excited by a light stimulation (presumably principal neurons). Expected stimulus onsets are indicated by black arrows. Shadings are 95% confidence intervals. Colored boxes above indicate the time courses of photo-stimuli. (**K**) Differences in the probability of stim-port licking by wS2 inhibition. Dots and bars correspond to each animal’s data and the average of all animals, respectively. Vertical lines on top of scatter and bar plots indicate 95% CIs. *: p<0.05 and <0.01, student’s t-test (null hypothesis: average=0). N=5 animals. (**L**) Differences in the probability of NoLick trials by wS2 inhibition. Conventions are the same as panel K. Significant difference from 0 was not observed in any condition. N = 5 animals.

There were five possible trial outcomes: (1) correct trials in which the stimulus was presented and the mouse indicated its presence by licking at the Stim port; (2) correct trials in which the stimulus was absent and the mouse licked at the NoStim port; (3) incorrect trials in which the stimulus was presented but the mouse licked to the NoStim port; (4) incorrect trials in which the stimulus was absent but the mouse licked to the Stim port; and (5) trials in which the mouse failed to respond with a lick at either port (“NoLick” trials). Correct trials were rewarded with a small drop of water delivered to the appropriate lickport. Other trial types were neither rewarded nor punished. After animals were well trained in the task, we titrated down the stimulus intensity so that mice made incorrect responses on ∼30% of trials (Figure 1D; Methods).

While mice performed the task, we used 64-channel silicon probe arrays to monitor spiking activity of single units in wS1 and wS2, in separate sessions (Figure 1C). Both wS1 and wS2 units showed more activity on trials in which mice made a lick response of any kind (whether to the Stim port or NoStim port, and for both Stim and NoStim trials) compared against activity on NoLick trials (Figure 1E,F; compare red and blue traces vs black traces). Thus, on average wS1 and wS2 neurons showed activity associated with licking per se, separate from any activity evoked by the tactile stimulus or associated with perceptual choice (Figure S2A-B).

When mice reported the presence of the stimulus by licking at the Stim port, activity in both wS1 and wS2 was greater on average compared with when they indicated stimulus absence by licking at the NoStim port (Figure 1E,F; compare blue vs red traces). Thus, units in both wS1 and wS2 also showed activity related to perceptual choice, rather than only to licking per se.

To quantify choice-related activity, for each unit we calculated a “choice probability” (CP)^6,7^. CP for a unit is the probability with which an ideal observer could predict a subject’s perceptual choice on a single trial---in our case whether the mouse responded at the Stim port or the NoStim port---based on the single-trial spiking activity of that unit. CP values calculated over [100, 500 ms] relative to the stimulus onset were on average significantly higher than chance level (= 0.5) for units in both wS1 and wS2 (Figure 1G). However, CP values in wS2 were higher than in wS1, for both Stim and NoStim trials (Figure 1G). Choice-related activity was observed among neurons throughout the cortical layers, with the exception of L4 in wS1 (Figure S1A-D). This lack of CP in wS1 L4 neurons may reflect the relative paucity of long-range cortico-cortical inputs to these neurons and the strength of inputs from VPM thalamus and other nearby L4 neurons^8–13^.

Together, these results show that activity in the somatosensory cortices of mice correlates both with perceptual choice and with licking per se, and that the association with choice is stronger in wS2 than in wS1.

### Inhibiting wS2 activity impairs perceptual choice but not licking

We used optogenetic excitation of ChR2-expressing parvalbumin (PV)-positive inhibitory neurons in wS2 to inhibit wS2 activity and thereby assay its role in task performance. We delivered light within one of two time windows: “early” ([0, 100 ms]) or “late” ([100, 500 ms]) relative to the time of possible whisker stimulus onset (Figure 1H-J) (see also: ^2^). The early window was chosen to cover the period with sensory responses to the whisker stimulus. The late window was chosen to cover sustained activity that could occur between stimulus offset and typical reaction times (Figure S1E-H). We carefully ramped down the light intensity to avoid a “rebound” of activity upon termination of the inhibition (Figure 1J)^14^. Inhibition during the early window biased mice toward reporting stimulus absence on Stim trials (Figure 1K). Inhibition during the late window also biased mice toward reporting stimulus absence, on both Stim and NoStim trials (Figure 1K). In contrast, inhibition of wS2 did not cause a significant change in the number of NoLick trials (Figure 1L). Together, these results suggest that wS2 activity is important for perceptual choice but not licking per se.

### Choice-related activity is enhanced when the sides of the stimulus and response are congruent

For the experiments reported thus far, animals were trained to report the stimulus via licking at the lickport located on the same side of the body as the stimulus (Right-stim→Right-lick). As animals’ sensory perception was measured by a specific action (Right-lick as opposed to Left-lick), we cannot tell if the choice related activity we observed was derived from the perceptual event, the stimulus-associated licking behavior per se or a combination of both. Therefore, we trained another group of mice (n = 5) with the opposite contingency between trial type and reward water port; Stim port on their left and NoStim port on their right (Right-stim→Left-lick contingency, Figure 2A, bottom). The former will be referred to as ‘congruent’ and the latter ‘incongruent’, with these terms referring to the correspondence between the side of the stimulus and the side of the port used to indicate stimulus presence.

**Figure 2.**
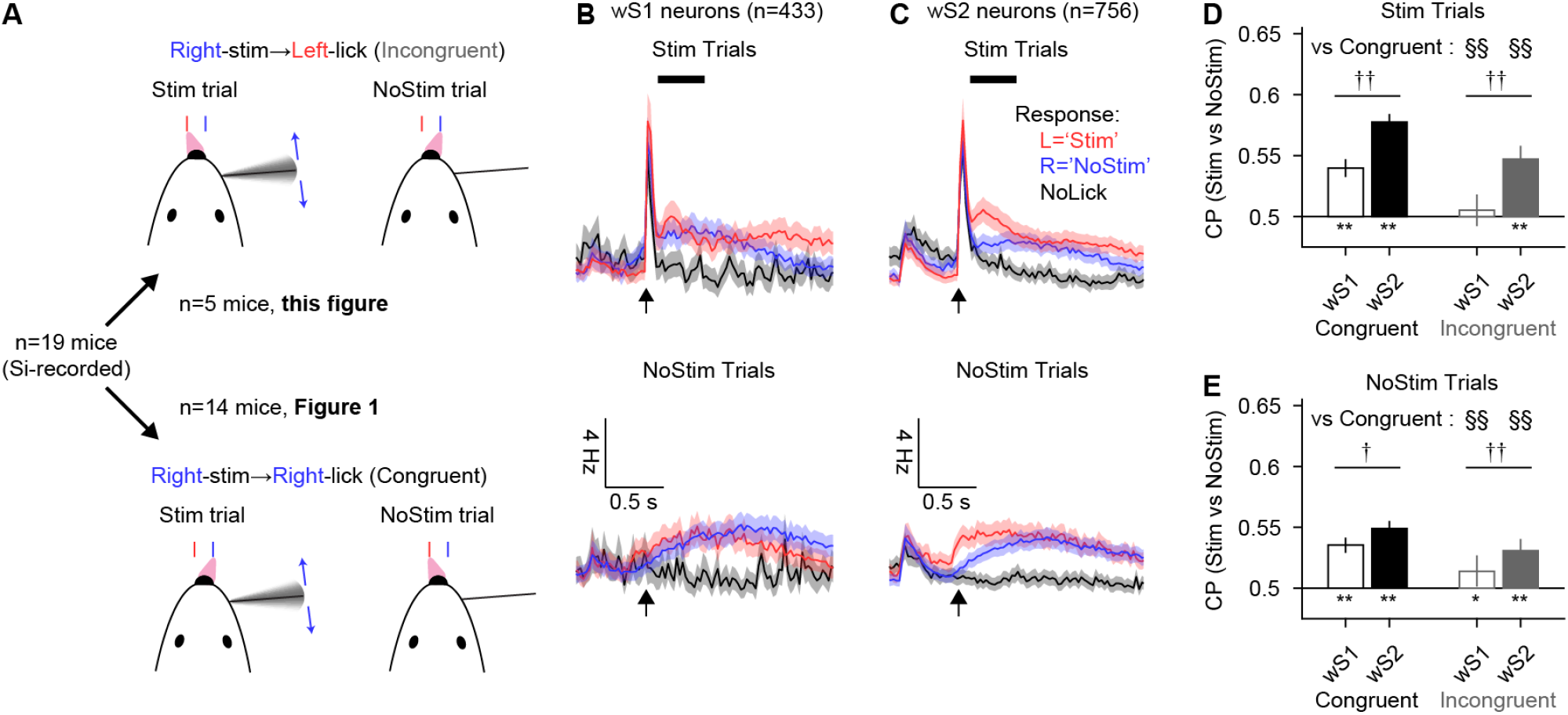
Stronger choice-related activity in congruent vs incongruent tasks. (**A**) A subset of animals were trained for a whisker detection task with the opposite contingency from Figure 1 (Right-stim→Left-lick, incongruent). (**B**-**C**) Spike rate of all silicon probe-recorded wS1 and wS2 neurons under ‘incongruent’ contingency task. 9 and 16 recordings in wS1 and wS2 from 5 animals. (**D**-**E**) Choice probabilities of Stim- over NoStim-port licking. Shadings are 95% CI. *, **: p<.05 and <.01, Student’s t-test (null hypothesis: CP=0.5); †, ††: p<.05 and <.01, Two-sample t-test (null hypothesis: CP_S1_ = CP_S2_). §, §§: p<.05 and 0.01, Two-sample t-test (null hypothesis: congruent = incongruent in the same neuron and trial types).

Neurons in wS1 and wS2 showed prominent licking-related activity in mice trained with the incongruent contingency (Figure 2B-C, Figure S2C-D; red and blue traces both exceed gray trace). In wS1, on average there was not a clear difference in activity associated with perceptual choice for the incongruent contingency (Figure 2B; red and blue traces overlap). Neurons in wS2 did, in contrast, show choice-related activity for the incongruent contingency (Figure 2C). Specifically, wS2 neurons showed greater mean activity on trials with Stim port licking compared with NoStim port licking (Figure 2C; red trace exceeds the blue trace). This difference in activity persisted for hundreds of milliseconds after stimulus onset (Figure 2C). As wS2 neurons robustly encoded animals’ choice in both contingencies, such activity is not solely derived from the required action (i.e., Left- versus Right-lick). However, choice-related activity was stronger with the congruent compared with the incongruent contingency, for both cortical areas and for both Stim and NoStim trials (Figure 2D-E).

As lick- and choice-related activity was observed on average across the population in the somatosensory cortex, irrespective of task contingencies, we assessed the degree to which individual single units signaled both perceptual choice and licking per se, as opposed to only one or the other. To do so, we performed ideal observer analysis to quantify how well a unit’s spiking on a given trial could be used to discriminate licking (“lick probability”, LP), and counted the number of neurons with statistically significant values for CP alone, for LP alone, or for both CP and LP (Figure S2B,D). Across both wS1 and wS2 in the congruent task, CP-alone units were most abundant (20% and 27% in wS1 and wS2, respectively), followed by LP-alone units (16% and 12%). Comparatively few units had significant values of both CP and LP (11% and 12% in wS1 and wS2, respectively). Thus, perceptual choice and licking per se were signaled by largely separate, but overlapping, populations of units. This was also true for the incongruent task, although CP-alone units were smaller in proportion.

Multiple studies have observed stronger choice-related activity among neurons that better discriminate the relevant sensory stimuli^3,6,15–19^. This would be expected if, intuitively, the brain relied on more discriminative neurons during perceptual choices. We quantified the ability of each neuron to discriminate between the presence vs absence of the whisker stimulus (“stimulus probability”, SP), and compared this quantity to a neuron’s choice probability. For the congruent task, we found modest correlations between CP and SP in both wS1 and wS2 (Figure S2E,G). However, these correlations were significantly weaker in the incongruent task (Figure S2F-G).

### Choice-related activity in S2→S1 neurons is enhanced for congruent responses

Next, we recorded from wS2 neurons that, based on optotagging, were classified as either S2→S1 neurons or S2→M2 neurons. We injected an AAV that expresses Cre-recombinase with a high efficiency of retrograde labeling (AAV-retro-Cre) into either wS1 or wM2. wM2 receives a dense wS2 projection and contributes to licking behavior in a whisker stimulus-based Go/NoGo discrimination task^20^. Another AAV with a Cre-dependent ChR2-expressing construct (AAV-DIO-ChR2) was injected to wS2 (Figure 3A,D). The former labels wS2 neurons that project to wS1 (S2→S1 neurons), and the latter S2→M2 neurons. As previously reported, S2→S1 neurons are densely localized in L4^21^ whereas histological and electrophysiological experiments showed that L4 was underrepresented among S2→M2 neurons, indicating non-overlapping distributions of those two groups of wS2 neurons (Figure S3). Larger spiking activity was evident in trials with Stim port licking under congruent contingency in both neuron types (Figure 3B,E, blue traces); that is, both populations of projection neurons showed significant choice probabilities. Consistent with the wS2 general population, both groups of projection neurons showed higher choice probability in the congruent task compared with the incongruent task (Figure 3G,H). Interestingly, choice-related activity was absent in S2→S1 neurons when animals were required to lick the port on the opposite side in response to a stimulus (incongruent). This sharp congruent-incongruent contrast is similar to that of wS1 neurons (Figure 2D,E), and may reflect a difference in their functional relevance between those two task types.

**Figure 3.**
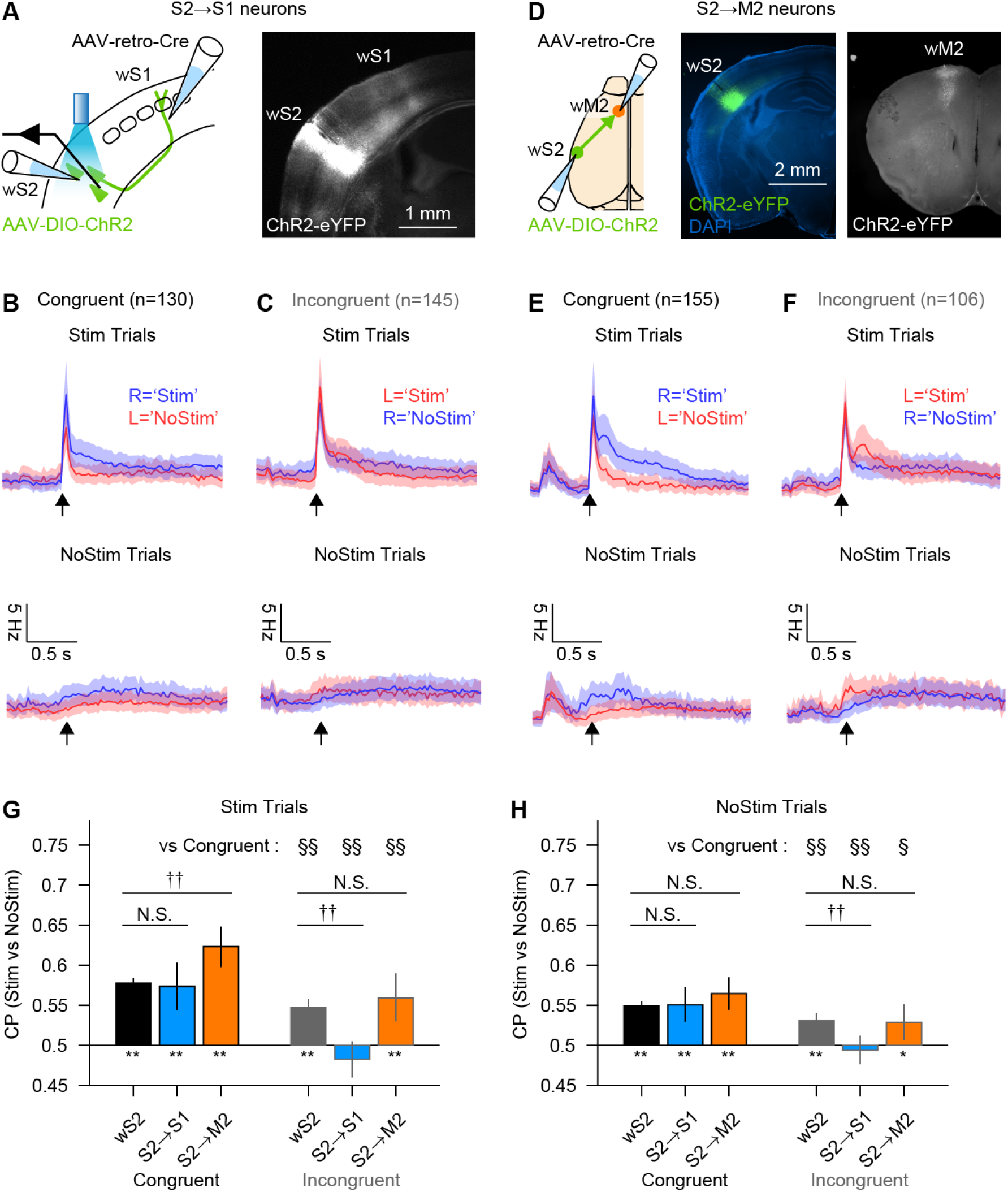
S2→S1 neurons exhibit choice-related activity specifically in the congruent task. (**A**) S2→S1 neurons were labeled with ChR2 by a combination of AAV-retro-Cre and AAV-DIO-ChR2 injections in wS1 and wS2 respectively. (**B**-**C**) PSTHs with 95% CI for S2→S1 neurons identified by a reliable response to photo-stimuli. Expected stimulus onset is indicated by an arrow. (**D**) Same as (A), but for S2→M2 neurons. Axonal projection in wM2 (right panel) indicates a successful labeling of wM2-projecting neurons. (**E**-**F**) Same as (B-C), but for S2→M2 neurons. (**G**-**H**) Choice probabilities of Stim- over NoStim-port licking. *, **: p<.05 and <.01, student’s t-test (null hypothesis: average=0.5); †, ††: p<.05 and <.01, two-sample t-test (null hypothesis: CP = CP_S2_ in the same neuron type). §, §§: p<.05 and 0.01, two-sample t-test (null hypothesis: congruent = incongruent in the same neuron type).

### Excitation of S2→S1 neurons promotes detection for congruent responses

Finally, we asked if excitation of those projection neurons affects behavioral performance. We optogenetically stimulated ChR2-expressing S2→S1 or S2→M2 neurons in the late response window ([100, 500 ms]). Units that were identified as ChR2-expressing cells (Methods) showed robust evoked activity in this time window (Figure 4A). Simultaneously recorded non-identified units were a mixture of excited, inhibited and non-affected units that on average showed a subtle change in activity (Figure 4B, Figure S4A-C). Activation of S2→S1 neurons robustly drove mice to lick at the Stim port during the congruent task, irrespective of trial type (Figure 4C). The same manipulation affected the licking response significantly less in the incongruent task. This striking contrast is consistent with the discrepancy in the strength of choice probabilities we observed when comparing S2→S1 neurons in the congruent vs incongruent tasks (Figure 3G,H). In contrast to these results with S2→S1 neurons, we found no effect of stimulating S2→M2 neurons, either for congruent or incongruent tasks (Figure 4D). This difference between S2→S1 neurons and S2→M2 neurons is consistent with functional differences between these major cortico-cortical outputs from wS2.

**Figure 4.**
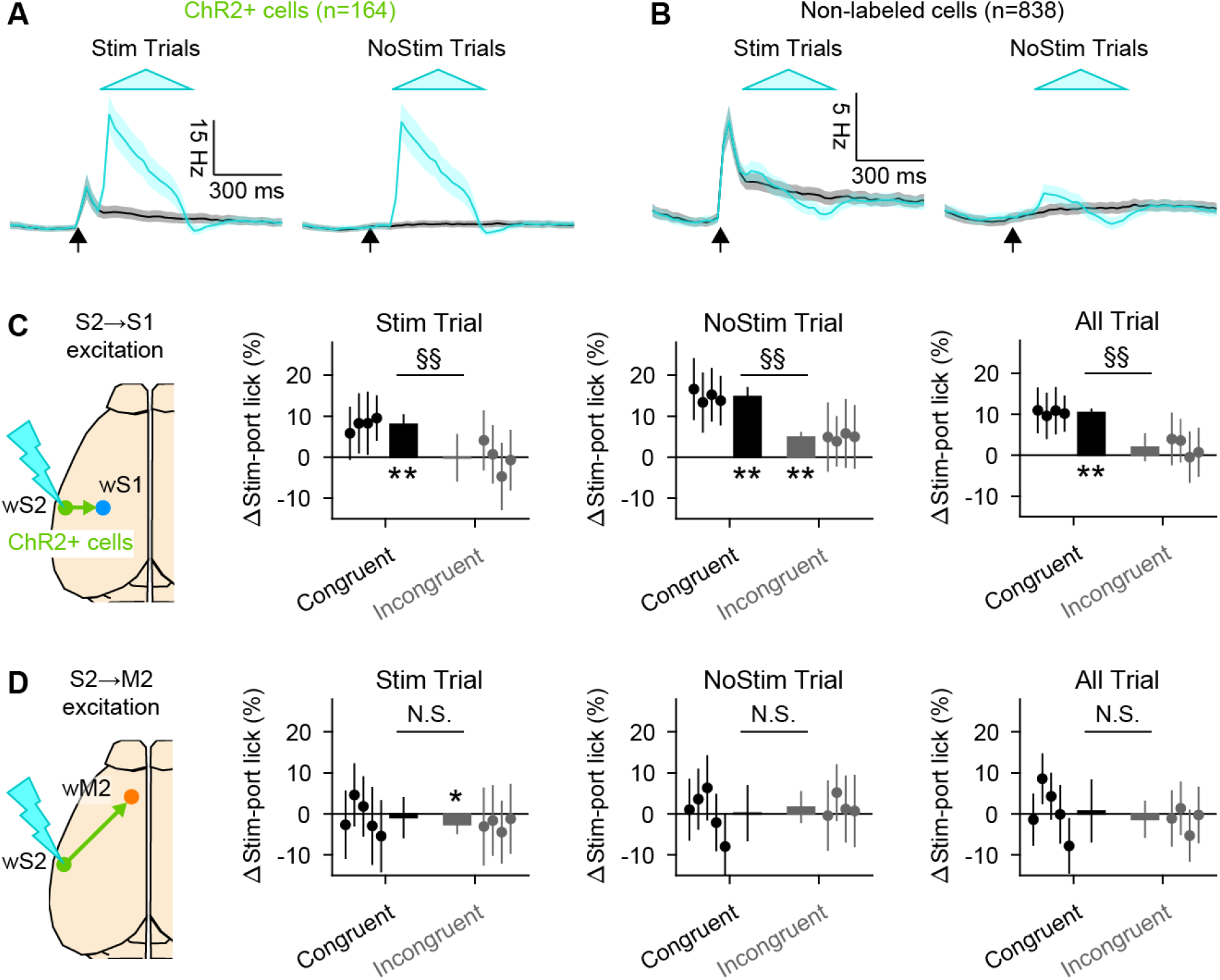
Excitation of S2→S1 neurons biases choice toward the stimulus-associated port in the congruent task. (**A**) PSTHs with 95% CI from identified ChR2+ cells in wS2 (i.e., S2→S1 and S2→M2 neurons). Cyan and black traces correspond to trials with and without a photostimulus. Cyan triangle indicates the timing and amplitude of photo stimulus. 19 sessions from 11 animals. (**B**) PSTHs of the cells that were not responsive to the photo stimulus in the same recording sessions as (A). (**C**) Behavioral effect of photo-excitation of S2→S1 neurons during the task. Differences in the probability of Stim-port licking by a photo stimulation for different task contingency (congruent/incongruent) and trial (Stim/NoStim/All) types. Each dot corresponds to each animal’s data, and a bar is the average of all animals in the same conditions. Error bars: 95% CI (for both scatter and bar plots). *, **: p<0.05 and <0.01, student’s t-test (null hypothesis: average=0). §§: p<0.01, two-sample t-test (null hypothesis: congruent = incongruent). (**D**) Same as (C) but for S2→M2 neurons.

## DISCUSSION

Here we have described multiple features of somatosensory cortex activity as it relates to performance of a licking-based perceptual detection task. Use of a task that required licking to report both the presence and the absence of a whisker deflection stimulus allowed us to distinguish neuronal activity associated with the perceptual choice from activity related to a licking response per se. This is generally difficult to achieve with Go/NoGo tasks in mice, where separating perceptual and licking signals requires a delay of hundreds of milliseconds or more to be interspersed between the offset of the stimulus and the time window in which a response is allowed^22,23^. While such tasks containing a delay-period offer a best-of-class method for isolating sensory and movement signals, they do not model the most common forms of sensorimotor interactions with touch. Normally, touch and movement exist in a tight closed-loop whereby sensory signals are used to guide ongoing and subsequent movements to interact with an object, as during whisker- or haptics-based exploration. In these cases, sensory and motor signals must to some extent overlap in time. Our intent was to investigate sensory and motor signals, and their potential interactions, in the whisker somatosensory cortex in this more typical case where they might occur simultaneously within the same population of neurons.

Our use of a two-alternative forced-choice task allowed us to observe distinct choice-related and lick-related activity in population means and in individual cells (Figures 1-2 and Figure S2). In our optogenetic inhibition experiments (Figure 1H-L), it also allowed us to observe an effect of wS2 inhibition on choice while ensuring that the optogenetic stimulus did not interfere with task-related licking nonspecifically. Best-of-class evidence for identifying decision signals in cortex, however, requires precise gain-of-function manipulations combined with neural and behavioral analysis^24,25^.

Choice probabilities in our experiments were higher among wS2 compared with wS1 neurons. This is consistent with prior work showing that activity in wS2 seems relatively more affected by the mouse’s perceptual report, as opposed to the stimulus, compared with wS1^1^. Similarly, systematic recordings from various brain areas during a vision-based motor task suggested a tendency that higher-order visual cortices display weaker sensory but stronger motor signals than primary visual cortex ^26^.

We observed choice-related activity across the cortical layers in wS1 and wS2, with the notable exception of layer 4 in wS1 (Figure S1). These L4 neurons project only locally in wS1, limited predominantly to their own barrel column, lack long-range corticocortical inputs, and receive their thalamic input from the ventroposterior medial (VPM) nucleus, a primary sensory nucleus. In contrast, L4 neurons in wS2 have long-range projections that target wS1^21^ and receive input from the posterior medial (POm) thalamic nucleus, a higher-order thalamic nucleus. Together, these observations suggest that choice signals reflect corticocortical or “top-down” activity^3,7,27^.

Choice-related activity was stronger in both wS1 and wS2 when the sides of the stimulus and response were congruent (Figure 2). This congruence effect of stimulus and action on sensory cortical activity seems to be task-type specific. While our recent study using a crossmodal sensory detection task also demonstrated a congruence effect in wS1^28^, other studies using more complex sensory discrimination tasks have not found such effects^25,29^. Because the connections linking somatosensory to motor cortices are predominantly ipsilateral (rather than contralateral), our data are consistent with a model in which choice-related activity in wS1 and wS2 propagates to ipsilateral motor areas to drive congruent (i.e. contraversive) licking. This model is also supported by our observation that excitation of S2→S1 neurons promoted only contraversive licking (Figure 4C). However, even in a Go/NoGo task without lateralized licking, sensory responses in mouse wM1 can be highly lateralized^30^. Preferential interactions among ipsilateral somatosensory and motor areas makes intuitive sense not only anatomically but also given the ubiquitous use of touch to interact with objects in the environment. Interestingly, in a simple reaction time task, humans showed better performance when they responded to a sensory stimulus with the same rather than the opposite side of the body^31^.

It is puzzling that while the choice probability was similar between S2→S1 and S2→M2 populations in the congruent condition, only photoactivation of the former had a behavioral effect. One explanation is that wS1 but not wM2 plays a pivotal role in linking sensation to action in a simple passive detection task such as ours, and choice-related activity in wS2 propagates to action-related areas largely by way of wS1. Our observation that inhibiting wM2 caused a small increase in responses to the Stim port does suggest some relevance of wM2 activity for our task (Figure S4), but it may be of greater importance in more complex tasks or those in which active sensing is required.

## METHODS

### Mice

All procedures were performed in accordance with protocols approved by the Johns Hopkins University Animal Care and Use Committee (protocol: MO18M187). We report anatomical experiments from 12 C57BL/6NHsd (Harlan) mice; 8 Ai9 (Jackson Labs: 007909; B6.Cg-Gt(ROSA)26Sor^tm9(CAG-tdTomato)Hze^/J) mice^32^; tetrode recordings in 9 C57BL/6NHsd mice from wS2; silicon probe recordings in 6 C57BL/6NHsd, 10 Ai9 and 5 PV-Cre mice from wS1 and wS2. All of 9 tetrode-recorded mice were male. 11 of 21 silicon probe-recorded mice were female. Ages ranged from 6 to 23 weeks at the start of experiments. Mice were housed in a vivarium with a reverse light-dark cycle (12 h each phase). Experiments occurred during the dark phase. Mice were singly housed after surgery and during behavioral experiments. Mouse cages included enrichment materials such as bedding and plastic domes (Innodome, Innovive).

### Adeno-associated viruses

We obtained rAAV5.EF1a-DIO-hChR2(H134R)-EYFP (“AAV-DIO-hChR2-EFGP”) from the UNC Gene Therapy Center Vector Core, and rAAV2-retro-Syn-Cre from the Janelia Virus Service Facility.

### Behavioral task

All behavioral experiments were conducted with head-fixed mice. Behavioral apparatus was controlled by BControl software (C. Brody, Princeton University). Seven to 10 days after surgery and 7-14 days before behavioral training, mice were allowed ∼1 ml of water daily until reaching ∼70% of their starting body weight. On training days, mice were allowed to perform until sated and were weighed before and after each session to determine the amount of water consumed. Additional water was given if mice consumed <0.3 ml of water. On the first day of training mice were acclimated to head fixation in the behavioral apparatus while being given free access to water via two reward ports located 6-10 mm and ∼35 degrees to the left and right of the mouse midline.

*Two port whisker detection task* On the subsequent days, animals were trained for detection task with a single whisker (always on the right whisker pad) threaded into a glass pipette attached to a piezo actuator (D220-A4-203YB, Piezo Systems), which was driven by a piezo controller (MDTC93B, Thorlabs). The whisker targeted at the time of ISI was used. Approximately 1.5 mm of whisker remained exposed at the base. All whiskers except the target whisker were trimmed to near the base. Every trial started with a 0.1 s auditory cue (8 kHz tone, ∼80 dB SPL) with a fixed 6.5-s interval. During all sessions, ambient white noise (cut off at 40 kHz, ∼80 dB SPL) was played through a separate speaker to mask any other potential auditory cues associated with movement of the piezo stimulator. 0.5 s after the onset of the auditory cue, a sinusoidal whisker deflection was applied in half of trials (Stim trials, the other half was NoStim trials). Animals were rewarded with a drop of water (∼3 μl) for licking the port associated with the trial type (Stim or NoStim) in an “answer period” spanning 0.3 s to 2 s after stimulus onset. Licks that occurred within a 0.3 s “grace period” immediately following stimulus onset were not counted.

Mice were trained for this two-port whisker detection task either with the “congruent” or “incongruent” contingency. In congruent contingency, the rewarding port in Stim trials (Stim port) was on the same side as the stimulus (i.e., right), whereas the rewarding port for NoStim trials (NoStim port) was on the other side (i.e. left). In incongruent contingency, the rewarding ports were reversed (i.e. Stim and NoStim ports were on the left and on the right, respectively). For the first ∼10 days, a 1-s whisker stimulus (sinusoidal, rostro-caudal deflections at 40 Hz, 1 s, ∼2000 deg / s) was used. Mice were manually given a drop of water from the correct rewarding port during the answer period after ∼5 trials without correct licks.

Performance was quantified as percent correct licks out of all the licks: 100*(#Correct licks) / (total # of trials with licks). After overall performance reached >80% without intervention, the stimulus amplitude and duration of 40 Hz whisker stimulation were reduced to ∼1000 deg/s and 0.025 s (i.e., 1 cycle) in the following ∼7 days, gradually enough so that the performance was maintained at the same level. Then a “censor period” ending at stimulus onset was introduced to facilitate correctly timed licking, duration of which was increased from 0.1 s to 0.25 s over the following ∼3 days. Premature licks occurring in this period resulted in the abortion of that trial with a 1 s alarming tone (white noise, cut off at 40 kHz, ∼100 dB SPL). Finally, the stimulus amplitude was reduced to a “near-threshold” level (300-800 deg/s varies from animal to animal) in which animals showed ∼70% overall performance over ∼5 days. Animals were regarded “well-trained” when they showed >50% performance for both Stim and NoStim trials with this near-threshold stimulus in 3 consecutive sessions, and moved onto manipulation/recording sessions.

Sessions in which overall performance was <60% correct, or <50% correct in either trial type, were omitted from further analysis. By these criteria, we excluded 11.9% of incongruent sessions (14 out of 118, from n = 11 mice) and 14.9% of congruent sessions (34 out of 228, from n = 18 mice).

### Headpost implantation

Titanium headposts were implanted for head fixation at 6-23 weeks of age as described^33^. Briefly, mice were anesthetized (1%–2% isoflurane in O_2_; Surgivet) and mounted in a stereotaxic apparatus (David Kopf Instruments). Body temperature was maintained with a thermal blanket (Harvard Apparatus). The scalp and periosteum over the dorsal surface of the skull were removed. The skull surface over the posterior half of the left hemisphere, which covers wS1 and wS2, was thinned with a dental drill. Unless otherwise stated, all the cortical areas mentioned (wS1, wS2 and wM2) are on the left hemisphere, which is contralateral to the whisker stimulated. The remaining exposed area of the skull was scored with a dental drill and the head post affixed using cyanoacrylate adhesive (Krazy Glue) followed by dental acrylic (Jet Repair Acrylic). An opening (“well”) in the head post over the left hemisphere was covered with silicone elastomer (Kwik-Cast, WPI) followed by a thin layer of dental acrylic.

### Intrinsic signal imaging

After recovery from headpost surgery (> 36 h), intrinsic signal imaging (ISI) was performed to identify wS1 and wS2 as described^21^. Briefly, mice were lightly anesthetized with isoflurane (0.5%–1%) and chlorprothixene (0.02 mL of 0.36 mg mL^–1^, intramuscular). In most cases, the target whisker was right D2. In rare cases D2 was missing at the time of ISI, and D1 or D3 was substituted. ISI was performed through the skull covered by a thin layer of cyanoacrylate adhesive. Since a noise from piezo actuator evokes activity in the vicinity of wS2, white-noise sound was played via a speaker placed near the animals.

### Virus injections

AAVs were injected via the thinned skull over the left wS1, wS2, wM2 using a beveled glass pipette (30–50 μm ID). wS1 and wS2 were localized by intrinsic signal imaging and injected at 4 different depths (from the pia: 300, 400, 600, 800 μm in wS1; 300, 500, 700, 900 μm in wS2) with the goal of covering the full thickness of cortex without spreading into the white matter. After the injection, the well in the head post was sealed with silicone elastomer and dental acrylic as described above. wM2 injections were conducted immediately prior to headpost attachment and were targeted by coordinates (AP: 1.8 mm, L: 1.0 mm) and injected with relatively larger volumes at 2 depths (100 nL at each depth at 500 and 1,000 μm). Thereby the injected area was covered by a layer of cyanoacrylate adhesive and by dental acrylic.

For retrograde labeling of neurons, rAAV2-retro-Syn-Cre was injected in the target area (wS1: 10 nL at each depth, 40 nL in total; wM2: 100 nL at each depth, 200 nL total at ¼ concentration) and AAV-DIO-hChR2-EFGP in wS2 (50 nL at each depth, 200 nL in total). For area-specific ChR2 expression in PV-Cre animals, a relatively smaller volume was injected in wS2 (20 nL at each depth, 80 nL in total) to limit the spread of virus. The period after injection till manipulations/recordings (recovery from surgery, water restriction and training) is about ∼6 weeks, and is enough for functional expression of ChR2.

### Optic fiber implantation

In a subset of animals, a 400-μm, 0.39 NA optic fiber was implanted above the wS2 or wM2 surface right after the AAV injection. A fiber tip on wS2 was covered with a small amount of eye-ointment which prevents silicon elastomer from blocking light penetration. The well of the headpost was covered with silicone elastomer and dental acrylic as in a normal headpost implantation. The optic fiber was fixed by a thin layer of dental acrylic but was easily detached for silicon probe recordings by drilling off the dental acrylic layer. A fiber implanted on wM2 was permanently fixed with cyanoacrylate adhesive and dental acrylic as a headpost was implanted.

### Tissue processing

Mice were perfused transcardially with 4% paraformaldehyde in 0.1 M PB (4% PFA) and brains were fixed in 4% PFA overnight (10-12 hr). Dorsal views of brains were imaged with stereo zoom microscope (Axio Zoom.V16, Zeiss) before those brains were sectioned coronally at a thickness of 80-100um. EYFP signal was amplified by immunostaining with chicken anti-GFP (1:500; A10262, ThermoFisher Scientific).

Sections were mounted on glass slides in DAPI-containing mounting medium (H-1200, Vector Laboratories). Images of processed tissues were acquired via CCD camera (QImaging, QIClick) and epifluorescence imaging (BX-41, Olympus; and Axio Zoom.V16, Zeiss).

### Electrophysiology

We recorded from multiple single units simultaneously using tetrodes or silicon probes in mice aged 10-30 weeks. Tetrode recordings were carried out using a custom-built screw-driven microdrive with eight implanted tetrodes (32 channels total)^34^. Tetrodes were implanted perpendicular to the cortical surface, which for wS1 and wS2 are ∼30 degrees and ∼55 degrees from vertical, respectively. Neural signals were filtered between 0.09 Hz and 7.6 kHz, and together with time stamps for whisker or optogenetic stimuli, were digitized and recorded continuously at 20 kHz (RHD2000 system with RHD2132 headstage, Intan Technologies). Spike waveforms were extracted by thresholding on bandpass filtered (700-6,000 Hz) signals and sorted offline using custom software. To measure unit isolation quality, we calculated the L-ratio^35^ and the fraction of inter-spike interval (ISI) violations for a 2 ms refractory period. All units included in the dataset had L-ratios < 0.05 and < 1% ISI violations. Because application of these two criteria require a sufficient number of spikes, we included only units with overall spike frequencies > 1 Hz, which corresponded to 3,000-5,000 spikes given the 60-80 min recording session durations (spikes from periods of repetitive photostimulation that began after the main recording session were not included). Recording sites were verified histologically using electrolytic lesions (15 s of 10 mA direct current).

Silicon probe recordings were via a 64-channel linear probe (ASSY-77 H3, version with sharpened tips, Cambridge NeuroTech). Recordings were done up to three times from an area with the same angle and depth but with slightly (50-100 μm) different positions in tangential plane. The probe was coated with DiI or DiD to histologically verify the recording area and to reconstruct the depths of channels within the cortex. The probe was inserted into the cortex at ∼30 degrees and ∼45 degrees from vertical for wS1 and wS2 recordings, respectively. The probe was left for 30 min before recordings for tissue relaxation. Animals were lightly anesthetized with 1-1.5% isoflurane prior to probe penetration and kept anesthetized until 15 min before the behavioral session. Neural signals were filtered between 0.09 Hz and 7.6 kHz, and together with time stamps for whisker or optogenetic stimuli, were digitized and recorded continuously at 30 kHz (RHD2000 system with RHD2164 headstage, Intan Technologies). Spike sorting was carried out with Kilosort^36^. We included neurons with < 1% ISI violations of a 2 ms refractory period. We excluded units that spiked on < 95% of the stimulus trials within a session, or with unstable spike shapes assessed by visual inspection. For reconstruction of channel positions within the cortex, the DiI- or DiD-marked point of the deepest probe tip insertion was located with respect to the bottom of the cortex. Additionally, we used whisker-evoked current source density (CSD; “CSDPlotter” package in MATLAB) analysis to estimate the middle of L4 for each recording.

Since silicon probe recordings were done with a linear probe penetrated at the center of ISI-targeted area, recordings were strictly confined to the representing areas of the whiskers we used for the task. Whereas tetrodes were implanted as a group of eight, and thereby some of them penetrated outside of the areas we aimed at. Therefore, we used silicon probe recorded data by referring to general wS1 and wS2 units. Tetrode-recorded units that were optogenetically identified were used as S2→S1 neurons, but were not included in the general wS2 population to avoid sampling bias. Those ChR2-labeled neurons can be regarded as specific whisker-representing wS2 neurons because i) S2→S1 projection is topographically organized, and ii) those wS1-projecting neurons were labeled by a confined injection of a retrograde virus into an area representing a specific whisker.

### Optogenetic identification

Neurons labeled with ChR2 according to their projection target were identified by repetitive photostimuli applied at the end of recording after the behavioral session. For tetrode recordings, a 200-μm, 0.39 NA optic fiber was co-implanted with the tetrode microdrive such that its tip was < 0.5 mm above the surface of the D2 and surrounding whisker-representing area (targeted by ISI) in wS2. The optic fiber was coupled to a 473 nm laser (DHOM-L-473-200mW, UltraLasers) with intensity controlled by an acousto-optic modulator (MTS110-A3-VIS, QuantaTech). In animals recorded with silicon probes, the tip of a 400 μm, 0.39 NA optic fiber (Thorlabs) was placed ∼2 mm above the surface of wS2. The optic fiber was coupled to a 470 nm LED (M470F3, Thorlabs).

After each recording session, trains of 20 light pulses (2 ms pulse duration, 5 mW from the fiber ending) were delivered at 20 Hz. These light pulse trains were repeated for 30 cycles (600 pulses total). ChR2-labeled neurons were identified by the reliability of the photo-evoked response. Spikes occurring at the determined latency ± 3 ms relative to the onset of each light pulse were counted separately for the first to the twentieth light pulse within a train (summing across all 30 repetitions of the stimulus pulse train). These 20 spike counts were compared with baseline activity. If a light-evoked spike count exceeded mean + 1.96 SD of these binned baseline activity values (bin size = 6 ms), the response was regarded to be “significant” for that particular light pulse within the 20-pulse sequence. Neurons that showed at least 18 significant responses out of the 20 total pulses were considered to respond reliably and assigned as ChR2-positive neurons. More detailed description is found in: ^21^.

### Optogenetic manipulation

We carried out optogenetic manipulation of neuronal activity while mice aged 12-30 weeks were engaged in behavioral task. In tetrode-recorded animals, 4-5 manipulation experiments were conducted after all recording sessions, whereas manipulation experiments were done ∼3 and ∼2 times prior to and simultaneously with silicon probe recordings for each condition.

For optogenetic excitation of ChR2-labeled projection neurons, 470 nm light was delivered through optic fibers mentioned above (200-μm ID for tetrode recordings and 400-μm for silicon probe recordings) in ∼33% of trials both in Stim and NoStim trials. The light delivery spanned [100, 500 ms] from the expected stimulus time with 200-ms ramp-up and ramp-down periods (i.e., there was not constant phase). The maximum light power was 4 mW. In order to prevent animals to be informed about the presence or absence of this stimulus, LED stimuli (∼10 mW, 5 ms at 20 Hz during [-100, 500 ms]) were delivered in every trial from another optic fiber left inside a protective cone that was used to protect the tetrode microdrives. The fiber tip aimed at the dental acrylic on the mouse head inside the cone.

For inhibition of cortical areas by excitation of ChR-labeled PV+ neurons, 40Hz sinusoidal photostimuli were applied at two windows ([-25, 75 ms] and [100, 500 ms]) with 100-ms ramp down to prevent rebound firing of excitatory neurons. The light was delivered with each condition in ∼25% of trials. The mean light power during the constant phase was 0.75 mW during recording. For manipulation experiments before recordings, light power was doubled (mean 3 mW) with the assumption that the intact skull would block ∼50% of light. ‘Masking’ lights (a 473 nm laser) were applied to the animals’ eyes from ∼7 cm away from a decoupled optic fiber (∼6 μW, 5 ms at 20 Hz during [-100, 500 ms]) in every trial.

### Data analysis: Electrophysiology

We combined data from tetrode recordings and from silicon probe recordings. Spike rates were reported for every 25 ms bins. A receiver operating characteristic (ROC) analysis was used to calculate choice probability (CP) of Stim port over NoStim port. Detailed descriptions are found elsewhere. Briefly, 25 ms-binned spike activity was calculated and trials were grouped according to the animals’ licking of the Stim and NoStim ports. Trials with premature licks or with no licking in the answer period were excluded from the analysis. For each time bin and for each neuron, the area under the ROC curve (AUC) which corresponds to the CP was calculated by using “perfcurve” function in MATLAB. Stimulus probability (SP) and lick probability (LP) were calculated likewise. Stim trials and NoStim trials were compared for SP regardless of the animals’ response. Trials with Stim and NoStim port licking were pooled as ‘Lick trials’ and were compared with NoLick trials to calculate LP.

### Data analysis: Behavioral effects of manipulation

We quantified the effects of optogenetic manipulations of neuronal activity by calculating the change in probability of licking Stim-port (ΔStim-port licking) compared to control trials without photo stimulation. Data from all manipulation experiments in an animal was pooled and the trials with premature licks or without licking in the answer period were excluded. Then, those licking trials were sorted according to the outcome (Stim- or NoStim-port licking) and photostimulation conditions. ΔStim-port licking for the animal in a particular photostimulation condition was calculated by the following equation.

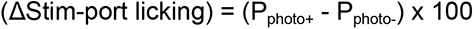

where P_photo+_ and P_photo-_ indicate proportions of Stim-port licking with and without photostimulus, respectively. Standard error (SE) of the difference in two proportions were calculated as follows:

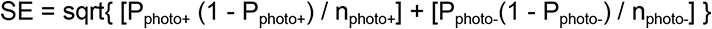

where n_photo+_ and n_photo-_ indicate the numbers of observations (trials) in each condition. By assuming z-distribution, 95% confidence interval was calculated;

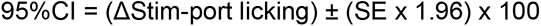

### Quantification and Statistical Analysis

Data analyses were conducted in MATLAB. Data are reported as mean ± 95% confidence intervals that were calculated using a nonparametric multistage bootstrap method unless otherwise stated. All statistical tests were two-tailed.

## Data and Code Availability

Data and MATLAB code used for analyses are available from the corresponding author upon request.

## Acknowledgements

We thank Angel Delgado, Julia Dursi, Lily Steele-Dadzie and Vincent Hou for technical assistance. This work was supported by NIH grants R01NS089652, 1R01NS104834-01 and P30NS050274.

---

**Figure S1.**
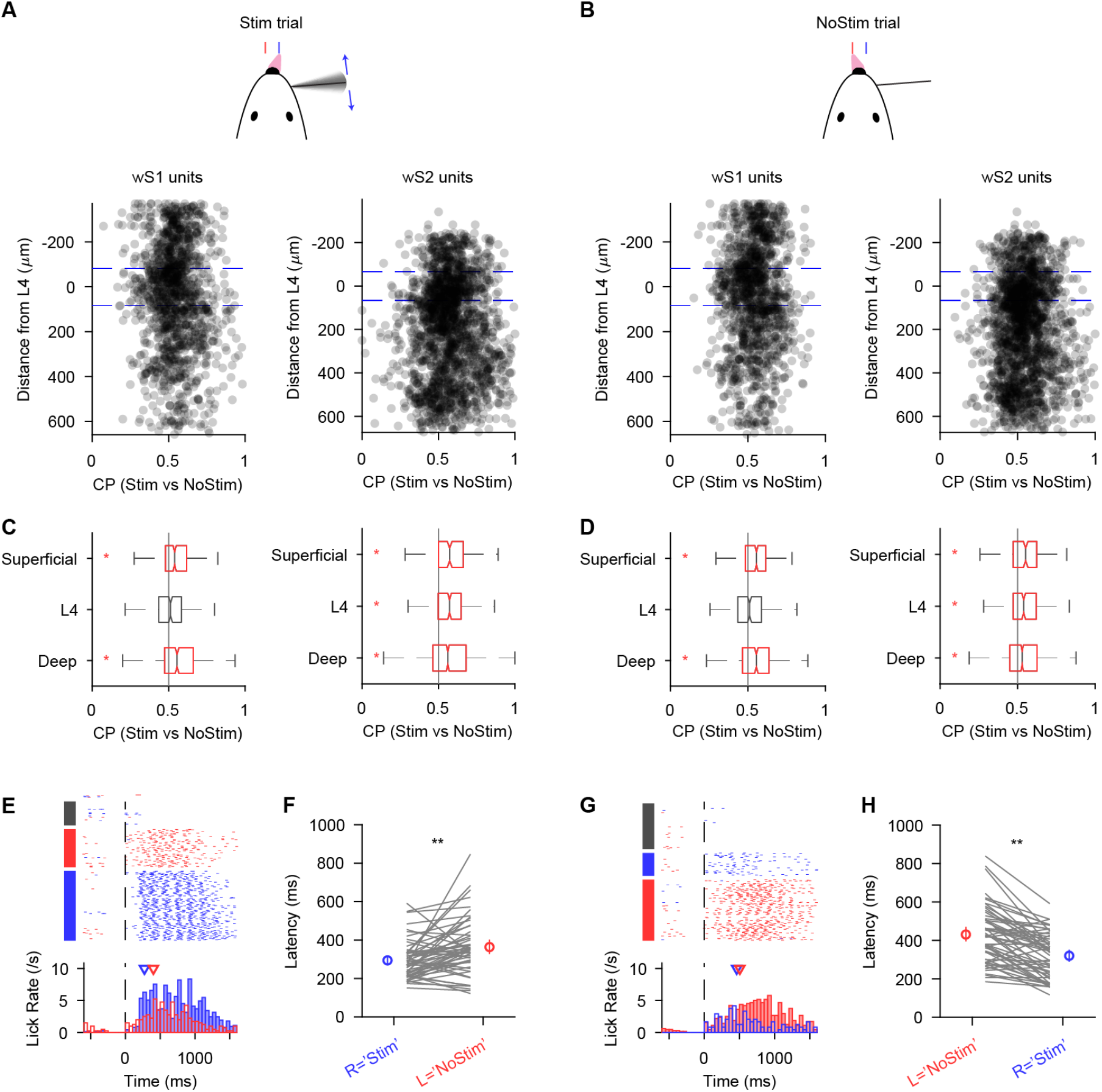
Choice-related activity is absent in L4 neurons of wS1. (**A-B**) Choice probabilities as a function of recorded depth of units. (**C-D**) Box plot of choice probabilities for different layer types. *: p<0.01, Wilcoxon signed rank test. (**E**) Lick behaviors of an example animal in a session during Stim trials. Top, lick raster plot sorted by trial outcomes. Bottom, lick rate histogram triggered by the expected stimulus onset. Lick directions are indicated by colors of the ticks, bars or boxes on the left (Left: red, Right: blue). (**F**) First lick latency. (**G-H**) Same as (E-F) but for NoStim trials.

**Figure S2.**
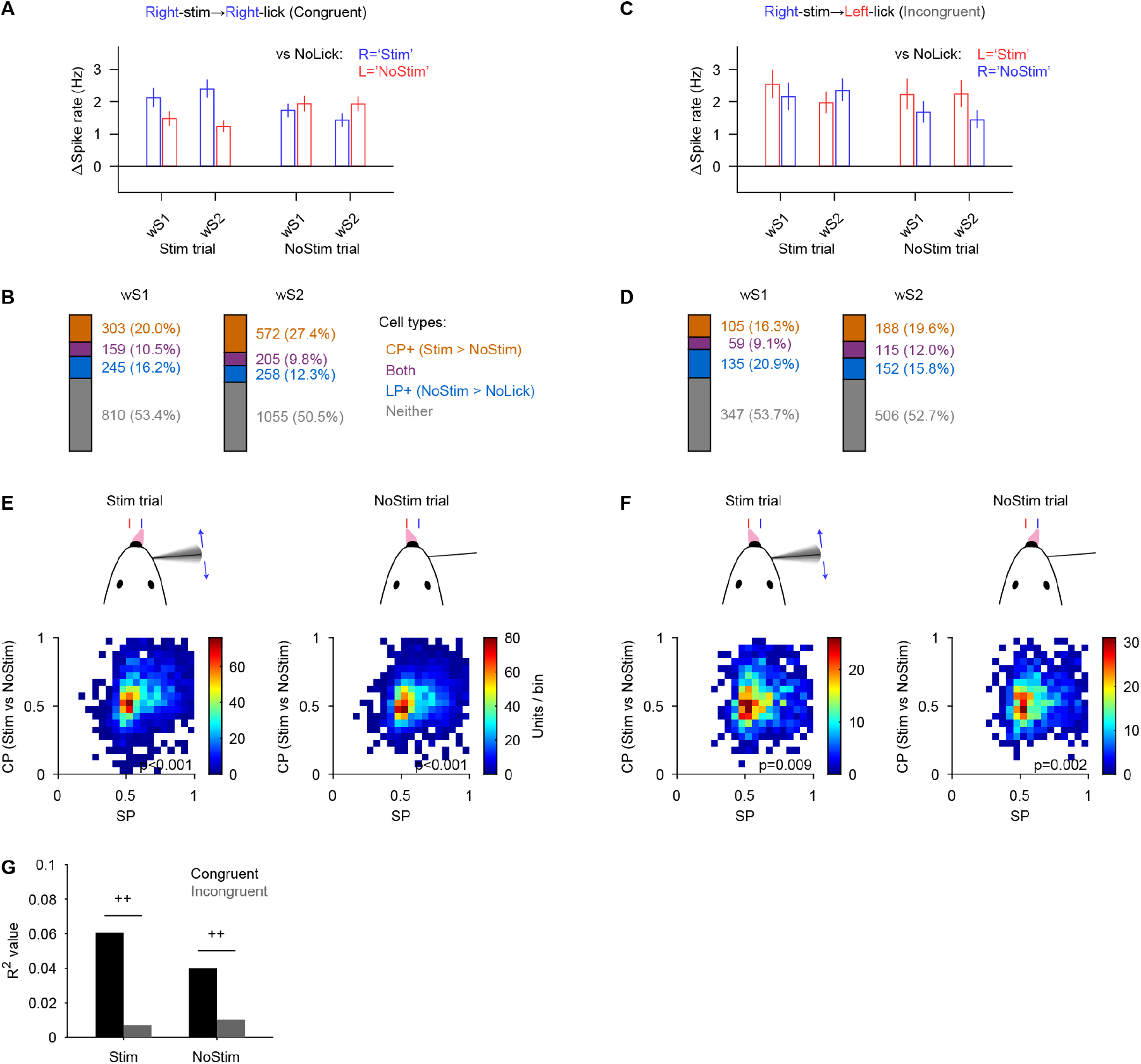
Somatosensory cortical neurons signal licking, choice and stimulus. (**A**) Spike rate difference of Lick from NoLick trials calculated for each condition at [100, 1500 ms] from the expected stimulus onset while animals were engaged in the congruent contingency task. Colors of bars indicate the side of the lickport. (**B**) Proportions of choice- and lick-reporting units (CP+ and LP+, respectively) in wS1 and wS2 during congruent task. (**C**) Distribution of choice probabilities (CP) as a function of stimulus probability (SP) for each trial type. (**D-F**) Same as (A-C) but for incongruent task. (**G**) Comparison of squared correlation coefficients (R2) between CP and SP. ++: p<0.01, Null hypothesis: z1=z2 after Fisher z-transformation.

**Figure S3.**
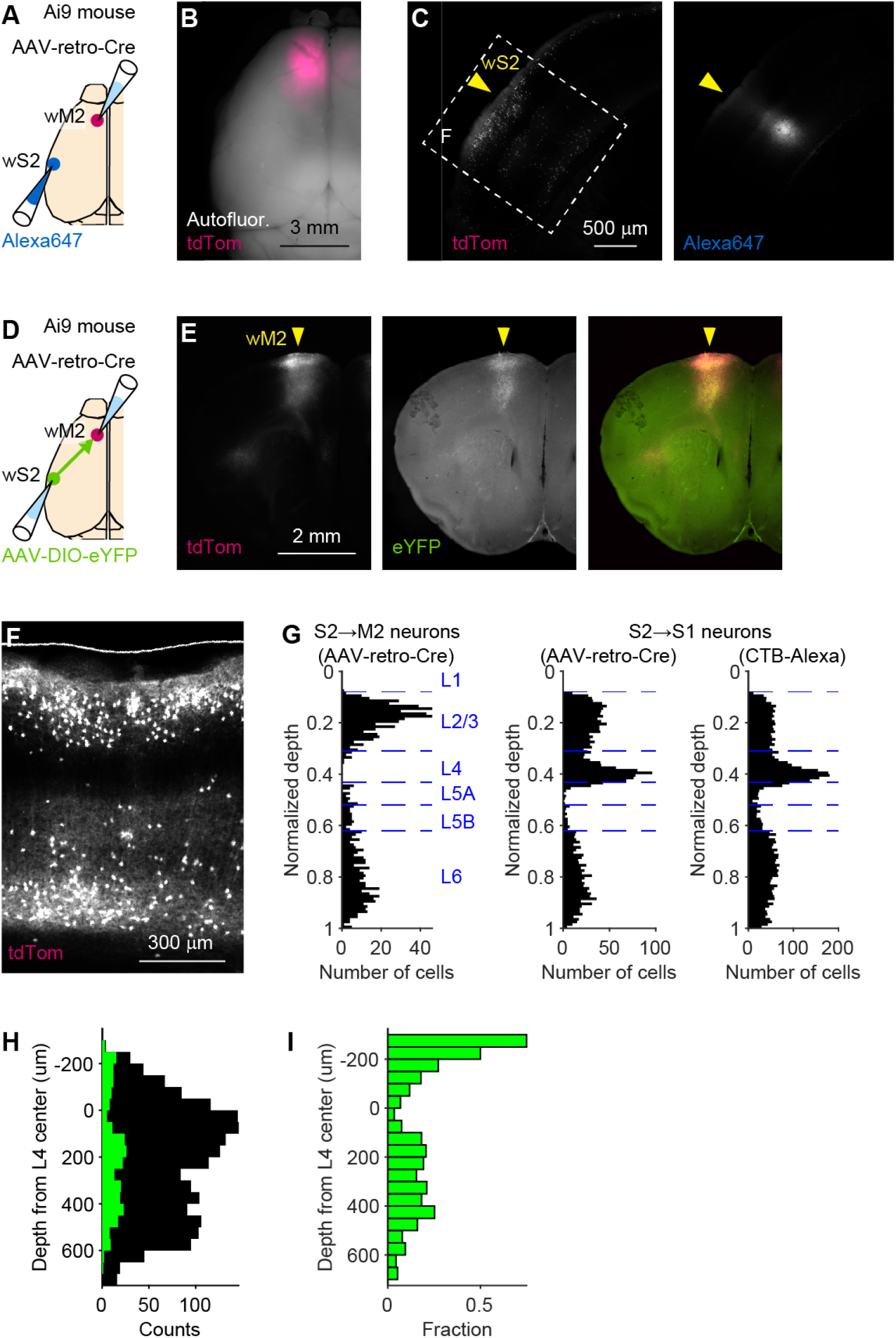
Viral labeling of S2→M2 neurons was confirmed by antero-/retro-grade tracing and by laminar distribution analysis. (**A**) AAV-retro-Cre and Alexa647 were injected to wM2 and wS2 of an Ai9 mouse. (**B**) Macroscope image of tdTomato and autofluorescence (green) were overlaid. (**C**) Retrogradely labeled S2→M2 neurons. Right, wS2 coordinate was confirmed by the presence of Alexa647 injected according to the ISI signal. (**D**) AAV-retro-Cre and AAV-DIO-eYFP were injected to wM2 and wS2 of an Ai9 mouse. (**E**) S2→M2 neurons showed strong projection in wM2. (**F**) An example epifluorescent image of retrogradely labeled S2→M2 neurons. (**G**) Laminar distribution of retrogradely labeled S2→M2 neurons (n = 6 sections from 6 animals), showing L4 is largely under-represented in comparison with S2→S1 neurons (data from Cell reports paper). (**H**) Laminar distribution of Si probe-recorded S2→M2 neurons (green) overlaid with all wS2 neurons (black) recorded from the same animals. (**I**) Fraction of S2→M2 neurons out of all recorded wS2 neurons in each depth. Note that under-representation of L4 is consistent with (G).

**Figure S4.**
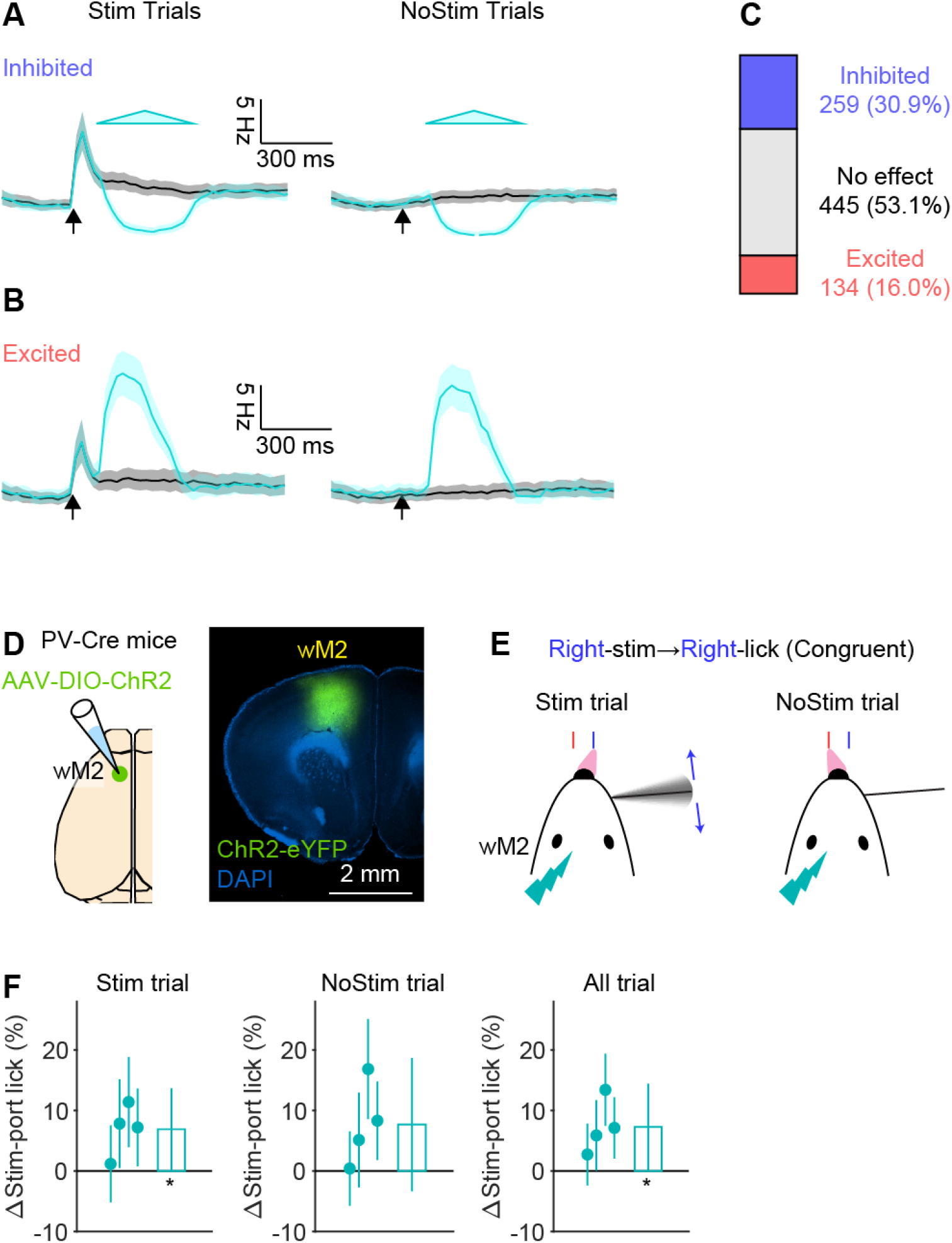
Mixed response of surrounding neurons by photo-excitation of a subset of wS2 neurons. (**A**) Mean spike activity of non-photo-identified units that were inhibited by photo-stimulation of ChR2+ units in Stim and NoStim trials, respectively. 95% CI is indicated by shading. (**B**) Same as (A) for units activated by photo-stimulation. (**C**) Proportion of units that were affected by photo-stimulation. (**D**) AAV-DIO-ChR2-eYFP was injected to wM2 of PV-Cre mice. (**E**) Schematic of whisker detection task in ‘congruent’ contingency with wM2 photo-stimulation. (**F**) Effect of wM2 inhibition on behavioral performance. From left to right, differences in the probability of Stim-port licking by a photo stimulation for Stim and NoStim trials, and for the entire session (All trial). Each dot corresponds to each animal’s data, and a bar indicates the average of all animals. Error bars: 95%CI. *: p<0.05 and <0.01, student’s t-test (null hypothesis: average=0). N=4 animals.

